# The Effect of Mechanical Stimuli on the Phenotypic Plasticity of Induced Pluripotent Stem Cell-Derived Vascular Smooth Muscle Cells in a 3D Hydrogel

**DOI:** 10.1101/2023.09.11.557280

**Authors:** Elana M. Meijer, Rachel Giles, Christian G.M. van Dijk, Ranganath Maringanti, Tamar B. Wissing, Ymke Appels, Ihsan Chrifi, Hanneke Crielaard, Marianne C. Verhaar, Anthal I.P.M. Smits, Caroline Cheng

## Abstract

**Introduction:** Vascular smooth muscle cells (VSMCs) play a pivotal role in vascular homeostasis, with dysregulation leading to vascular complications. Human induced pluripotent stem cell (hiPSC)-derived VSMCs offer prospects for personalized disease modeling and regenerative strategies. Current research lacks comparative studies on the impact of 3D substrate properties under cyclic strain on phenotype adaptation in hiPSC-derived VSMCs. Here we investigated the potential of human mural cells derived from hiPSC-derived organoids (ODMCs) to undergo phenotypical adaptation under various biological and 3D mechanical stimuli.

**Methods and results:** ODMCs were cultured in 2D conditions with synthetic or contractile differentiation medium, or 3D Gelatin Methacryloyl (GelMa) substrates with varying degrees of functionalization and percentages to modulate material stiffness, elasticity, and crosslink density. Cells in 3D substrates were exposed to cyclic unidirectional strain. Phenotype characterization was conducted using specific markers through immunofluorescence and gene expression analysis. Under static 2D culture, ODMCs derived from hiPSCs exhibited a VSMC phenotype, expressing key mural markers, and demonstrated a level of phenotypic plasticity like primary human vSMCs. In static 3D culture, higher substrate stiffness, lower elasticity and higher crosslink density promoted a contractile phenotype in ODMCs and vSMCs. Dynamic stimulation in 3D substrate promoted a switch towards a contractile phenotype in both cell types.

**Conclusion:** Our study demonstrates a phenotypic plasticity of human ODMCs in response to 2D biological and 3D mechanical stimuli that equals that of primary human vSMCs. These findings may contribute to the advancement of tailored approaches for vascular disease modelling and regenerative strategies

## Introduction

Vascular smooth muscle cells (vSMCs) provide structural vascular support and stability, regulate blood flow, contribute to immune responses and participate in tissue repair mechanisms ^1–3^. Dysfunctions in vSMCs lead to vascular complications with significant impact on morbidity and mortality, including (non-)constructive coronary artery disease (CAD), carotid artery and aortic aneurysms and pulmonary arterial hypertension ^4–8^. Understanding the cellular processes that contribute to disease etiology is essential for developing dedicated *in vitro* disease models, targeted therapies as well as regenerative strategies for the treatment of vascular diseases.

Following differentiation during vascular development, vSMCs retain a considerable degree of plasticity, manifested in a spectrum of phenotypic variations that encompasses synthetic characteristics involving proliferation, extracellular matrix (ECM) synthesis, and tissue repair, as well as contractile properties involving force generation ^6, 9, 10^. During neovascularization, vSMCs exhibit a synthetic phenotype characterized by high rates of proliferation, cell migration, and deposition of ECM ^3, 4^. In mature functional blood vessels, vSMCs typically assume a contractile phenotype to fulfil an essential role in vessel stabilization and vasomotion, displaying a quiescent state with an elongated, spindle-shaped morphology ^4^. Phenotypic switching between contractile to synthetic occurs during adulthood in response to e.g. vessel injury and represents a critical step in the repair process ^5^.

Although animal models and primary vSMC culture systems have provided valuable insights into vascular biology and disease, there are substantial disparities in the vascular physiology between rodents and humans. In addition, limited availability of patient-derived vSMCs and the absence of well-characterized patient-specific three-dimensional (3D) tissue models that could better mimic the dynamic *in vivo* environment, currently impede advancements in research. Human-induced pluripotent stem cell (iPSC)-derived vSMCs offer an alternative platform for studying human vascular biology ^11–17^. Generated from healthy donor or patient-derived somatic cells, iPSCs derived tissue cells represent an abundant source for disease modeling, drug screening, and tissue engineering. Concerning vSMCs derived from iPSCs, several significant challenges remain to be addressed. One key issue is the phenotypic and developmental heterogeneity observed in many culture protocols, resulting in a mixed population of vSMCs with varying levels of synthetic and contractile phenotypes, as well as a degree of “contamination” of the cell pool with non-vSMCs. Several strategies have been developed for enrichment of lineage-or phenotype specific vSMCs while excluding non-vSMCs ^18, 19^. Yet most published studies have still relied on differentiated vSMCs with unclear embryonic origin, purity, or functional phenotype. This phenotypic heterogeneity in particular poses a significant challenge in their application for human disease modeling as well as regenerative medicine: The successful replication of physiological function relies on the presence of contractile vSMCs, whereas in vascular diseases as well as tissue (re)generation, synthetic vSMCs play critical roles. This duality becomes particularly problematic as iPSC-derived vSMCs therefore need to effectively mimic either contractile and synthetic phenotypes to accurately represent various disease conditions *in vitro* or archive the desired cellular phenotypes in different phases of vascular tissue engineering. An improved understanding of how environmental factors define (h)iPSC-derived vSMC phenotypes, could provide leads to possible solutions.

Various biological (growth) factors are known to induce the phenotypic switch in vSMCs *in vitro*, including Platelet Derived Growth factor B (PDGFB), that can be used to induce synthetic characteristics in primary VSMCs ^18, 20^, and transforming growth factor beta (TGF-β), which is reported to reverse synthetic VSMCs into a contractile phenotype ^19^. In addition, phenotype transformation of vSMCs is known to be modulated by mechanical cyclic strain, as shown in various *in vitro* studies, mimicking *in vivo* vascular dynamics. VSMC responses are variable, depending on the applied strain frequencies, elongation, and orientation (e.g. uni or bidirectional, on a flat or circumferential surface) ^21–23^. Phenotype determination in primary vSMCs may also be controlled by intrinsic matrix properties like stiffness ^24, 25^ and cell responses appear to significantly differ between 2D (cell grown on top of matrix substrate) versus 3D (grown in matrix substrate) culture conditions ^26–28^.

For (h)iPSC-derived vascular smooth muscle cells (vSMCs), previous studies have demonstrated that both biological growth factors and dynamic strain can initiate phenotypic adaptation, resembling the response of primary vSMCs ^14, 22^. Nevertheless, the influence of intrinsic matrix substrate properties, such as hydrogel stiffness, elasticity, and degree of crosslinking on (h)iPSC-derived vSMCs in especially 3D instead of 2D structures, with 3D more closely mimic physiological conditions, has not yet been investigated. Moreover, the phenotypic adaptation of (h)iPSC-derived vSMCs in response to these factors in 3D environments under cyclic strain, remains unexplored.

In this study, we evaluated the potential of human mural cells derived from hiPSCs obtained from vascular organoids (organoid derived mural cells, or ODMCs) to adapt into vSMCs with a contractile or synthetic phenotype. In particular, their response to various phenotype differentiation inducers, such as biological growth factors (TGF-β and PDGFB) are evaluated. In addition, we will assess their phenotypic responses to differences in matrix properties (stiffness/elasticity and crosslinking) in a 3D environment, using Gelatin Methacryloyl (GelMa) with different degrees of functionalization (DOF) and weight percentages, in the presence and absence of cyclic unidirectional strain. By investigating the impact of these environmental components on the phenotypic switch in ODMCs, the optimal conditions can be defined to grow and maintain a hiPSCs-derived vSMC pool with the desired phenotype. The findings from this study thus provide valuable strategies for complex *in vitro* modeling of vascular diseases, and have implications for regenerative approaches.

## Methods

### 1. 2D Growth-factor experiments

#### 1.1 2D Cell Culture

ODMCs were differentiated and harvested from hiPSC-derived blood vessel organoids as described previously ^29^. Aortic VSMCs were purchased from Lonza. Both ODMCs and VSMCs were cultured on 1% gelatin coated cell culture plates in SMGM2 medium (Lonza) at 5% CO_2_. Medium was changed every other day. Cells were passaged for expansion or harvesting using Trypsin/EDTA (Gibco). Cells were use until passage 7.

#### 1.2 Growth-factor induced phenotypic switch

Cells were plated onto gelatin-coated 18mm coverslips (staining) or gelatin coated 6-well plates (PrestoBlue and gene-expression analysis). They were serum-starved (0.5% Fetal Bovine Serum (FBS) in DMEM) for 24 h before inducing the phenotypic switch. Control groups were cultured in DMEM with 10% FBS and 1% Pen/Strep (P/S). Synthetic groups were cultured in DMEM with 10% FBS, 1% P/S, 10ng/ml PDGF and 1ng/ml TGFβ. Contractile groups were cultured in DMEM with 0.5% FBS, 1% P/S and 1ng/ml TGFβ. All were kept at 37°C with 5% CO_2._

#### 1.3 PrestoBlue viability assay

Cells (*n* = 6 different vials per cell type) were seeded on a gelatin coated six-well plate with a cell density of 5*10^4^ cells per well. Cell viability was measured 24, 72 and 144h after growth factor treatment using PrestoBlue Cell Viability Reagent (ThermoScientific) according to manufacturer’s protocol.

#### 1.4 FACS analysis ODMCs

ODMCs were cultured and harvested using Trypsin/EDTA as described above. Cells were distributed in 96-well plate (25.000 cells per well) and subsequently stained with anti-CD31 and anti-CD140b antibodies (Table S1), together with Sytox blue (Invitrogen) to exclude dead cells. CytoFLEX flow cytometer (Beckman Coulter) was used for cell analysis and data analysis was performed using FlowJo software (Version 10.2).

### 2. 3D Static GelMa experiments

#### 2.1 GelMa hydrogel preparation

Two GelMa stocks with different Degree Of Functionalization (DOF) were prepared. For both; 10g type A gelatin from porcine skin (Sigma Aldrich) was dissolved in 100mL PBS at 60°C to obtain a 10% gelatin solution. Gelatin was modified with methacryloyl groups by addition of 8.75mL (80 DOF) or 5.45mL (50 DOF) methacrylic anhydride (Sigma) dropwise to the 100 mL gelatin solution. After 3 h, 400mL PBS was added and the solution was dialyzed against distilled water to remove salts and methacrylic acid for 7 consecutive days. Finally, the solution was lyophilized and stored at −80°C until further use.

To prepare hydrogels; radical crosslinking of solubilized GelMa in PBS was conducted in the presence of a photo-initiator. For this, a 0.1% 2-Hydroxy-2-methylpropiophenone photo-initiator (PI) (Irgacure, Sigma Aldrich^®^) was prepared using PBS. Lyophilized GelMa (5 or 10% w/v) was mixed with 0.1% PBS-PI and incubated for 15 min at 80°C to dissolve.

#### 2.1 GelMa swelling assay

30μL of pre-polymer solution of both 80 DOF and 50 DOF (5% and 10%) was pipetted on a petri dish between two spacers with a height of 0.45 mm and was covered with a sterile glass slide. The pre-polymer solution was placed under a 450 mW UV-light (OmniCure Series 2000, Excelitas) for 50 seconds. The hydrogel was removed from the glass plate and washed with PBS. Empty gels were incubated in PBS at 37°C for 24 h before mechanical testing and hydrogel swelling analysis.

Swollen GelMa hydrogels (3 per experiment for 5 different experiments) were weighed (ww) and subsequently dried by lyophilization. After that, dried weight (wd) of GelMa gels was obtained and the mass-swelling ratio (q) was calculated as q = ww/wd.

#### 2.2 Dynamic mechanical analysis - DMA

The Q800 Dynamic Mechanical Analyzer (DMA) (TA Instruments, Inc.) was used to test the hydrogel mechanical properties through a controlled force. The hydrogels were placed between the parallel-plate compression. A ramp force was applied at 0.010 N/min to 0.500 N with a preload force of 0.0010 N for 10 minutes. The Young’s Modulus was calculated by the slope of the most linear part (8 datapoints) of the stress-strain curve.

#### 2.3 Cell-laden 3D static GelMa hydrogels

The ODMCs/VSMCs were added to the GelMa-PI solution to achieve a cell density of 75 000 cells per 30 μL. The cell containing GelMa solution (30 μL) was pipetted on a 10 cm dish between to spacers covered with a sterile microscope slide and placed under a 450 mW UV-light (OmniCure Series 2000, Excelitas) for 50 seconds. The hydrogels were subjected to serum starvation (0.5% DMEM) for 24 hours after which 10% DMEM was added. *N*=6 per condition.

#### 2.4 Live/dead cell viability assay

A live/dead assay on the cell-laden GelMa constructs (*n=*6 per condition) was performed using a LIVE/DEAD^TM^ Cell Imaging kit (Invitrogen, Waltham, Massachusetts, United States) 144 h after Gel synthesis. The fluorescent dyes were diluted according to manufacturer’s instructions, in DMEM supplemented with 10% FBS and 1% P/S and added to the gels to incubate for 10 minutes at room temperature in the dark. The constructs were analyzed with fluorescence imaging on 470 nm and 550 nm wavelengths for the green and the red signal respectively.

### 3. 3D Dynamic experiments

#### 3.1 Cell-laden 3D GelMa Hydrogels exposed to dynamic loading

Two 5 x 20 mm Velcro strips were glued in parallel, 5 mm apart, to the bottom of each well of 6-well Bioflex culture plates (untreated, Flexcell Int) using medical adhesive silicone (Silastic MDX4-4210, Dow Corning, Midland, MI). Each pair of Velcro strips served as a mold to attach a hydrogel to the flexcell membrane. Cell-GelMa suspensions (750 000 cells per 300 μL each) were pipetted within these molds and placed under a 450 mW UV-light for 100 seconds. The hydrogels were covered with SMGM-2 medium and cultured for two days before dynamic loading was applied. At day 3, cell-laden GelMa hydrogels were placed on the Flexcell FX-5000T (Flexcell Int, McKeesport, PA) and exposed to 2 days of 10% strain (0.5 Hz). *N*= 3 per condition.

#### 3.2 Strain validation

To validate the intra and inter-experimental variations in dynamic loading, 3 x 3 dotted patterns were created on the membranes of a 6-well Bioflex culture plate (untreated, Flexcell Int). Videos were captured at day 3 (the first day of straining) and day 5. Subsequently, the maximum strain in the y direction (ɛ_yy,max_) was calculated (Supplementary fig. 3) by tracking the displacements of the previously applied dotted patterns over time using the open source software Tracker (https://physlets.org/tracker).

To validate whether hydrogel intrinsic properties affected the straining of cell-laden 3D GelMa hydrogels, 50-5 and 80-10 gels (n=6/group) without cells were created, covered with graphite particles, and imaged while being exposed to the 10% straining protocol. To assess whether cell remodeling activities would affect the strain pattern in the hydrogels, also strain patterns of cell-laden 3D GelMa hydrogels were assessed at day 5 in a similar fashion. The captured videos were converted to images at 30 Hz in MATLAB (Mathworks, Massachusetts, USA). Subsequently, the maximum strains in the y direction (ɛ_yy,max_) per loading regime group were calculated (Supplementary fig. 3) using the open source 2D DIC software Ncorr (v1.2, www.ncorr.com).

### 4. Analysis and Immunohistochemistry

#### 4.1 Quantitative Polymerase Chain Reaction Analysis

Total RNA was isolated from cultures (ODMCs and VSMCs) using RNA isolation kit (Bioline) according to the manufacturer’s protocol. Cells from 3D GelMa constructs were extracted using the QIAshredder columns according to manufacturer’s instructions. Supernatant was subsequently used for RNA extraction with the RNA isolation kit from Bioline as described above. The purity and concentrations of RNA were quantified using spectrophotometry (DS-11; DeNovix) absorbance measurements at 260/280 nm. cDNA synthesis was performed according to the instructions from the Bioline cDNA synthesis kit. Gene expression was determined using FastStart SYBR-green (Roche) following the quantitative polymerase chain reaction (qPCR) program: 8,5′ 95 °C, 38 cycles (15′′ 95 °C; 45′′ 60 °C) 1′ 95 °C, 1′ 65 °C, 62 cycles (10′′ 65 °C + 0.5 °C) in the SYBR-Green-Cycler IQ5 detection protocol (Biorad CFX384), performed in 384-well plates (Merck). The primer sequences used are listed in Table **S2**. All results were normalized for house-keeping genes ROPL and RPLP0, resulting in relative mRNA expression. In dynamic experiments, results were compared to static controls and represented as fold change (ΔΔCt).

#### 4.2 2D Immunohistochemistry

Phenotypic switch was induced in cells cultured on 18mm coverslips as described in 1.2. Cells were fixated after 24 h and 72 h using 4% PFA for 20 minutes. Cells were blocked using a 2% PBS/bovine serum albumin (BSA) solution for 30 min. The cells were stained with anti-Calponin overnight at 4 °C (Table **S3**). Thereafter, the staining solution was removed and the coverslips were washed three times with PBS. Secondary antibody incubation together with phalloidin was performed for 1 h at RT (Table **S3**). The coverslips were washed with PBS and counterstained with DAPI for 5 min. Coverslips were mounted on microscope glass with Mowiol 4–88. Samples were stored at 4 °C prior to imaging.

#### 4.3 3D immunohistochemistry

Cell-laden GelMa constructs were fixated using 4% PFA for 1hr at RT. Constructs were blocked and permeabilized using 3% FBS, 1% BSA, 0.5% Triton x-100 and 0.5% Tween in PBS for 2 h at RT. GelMa constructs were stained with anti-Smooth Muscle Actin α and anti-Calponin (Table **S3**) for 2 h at RT. The cells were washed three times with PBS-/Tween and secondary antibody (Table **S3**) incubation was performed for 2 h at RT. DAPI was used as a counterstain and GelMa constructs were mounted with Mowiol4-88. Samples were stored at 4 °C prior to imaging.

#### 4.4 Imaging and Analysis

Imaging was performed using the Leica Confocal SP8× (2D cultures; 63x magnifications) and the Leica Thunder microscope (10×, 20× and 40× magnifications for 3D GelMa constructs). Images were analyzed using ImageJ software (V1.47). 3D images were composed in LASX (version 3.5.7.23225).

#### 4.5 Statistical Analysis

The statistical analyses were performed using Graphpad Prism (version 8.3). Values are shown as individual data points with mean ± SEM. Prior to statistical testing, outliers were removed from the results when detected using a Grubbs’ test (alpha = 0.05). The paired, two-sided t-test and the ordinary one-way ANOVA test with Tukey post hoc test were used when appropriate. Experiments were performed at least in triplicate. The detailed sample size for each result is listed in the legend of the figures. A p-value of *p* ≤ 0.05 was accepted as statistically significant. Significance is further described in figure legends and results section.

## Results

### ODMCs are capable of growth-factor induced phenotype switching similar to primary human aorta derived VSMCs

ODMCs were harvested from human iPSC-derived blood vessel organoids following a previously described protocol summarized in figure 1A. Blood vessel organoids, cultured following the adapted Wimmer lab protocol ^29, 30^ contain both endothelial (CD31+) and mural (CD140b+) cells in a vasculature-like organization (fig. 1B). Extracted ODMCs bought into single culture express both αSMA and CD140b (fig. 1C-E) and were devoid of endothelial cells contamination as confirmed by FACS analysis (fig. 1E).

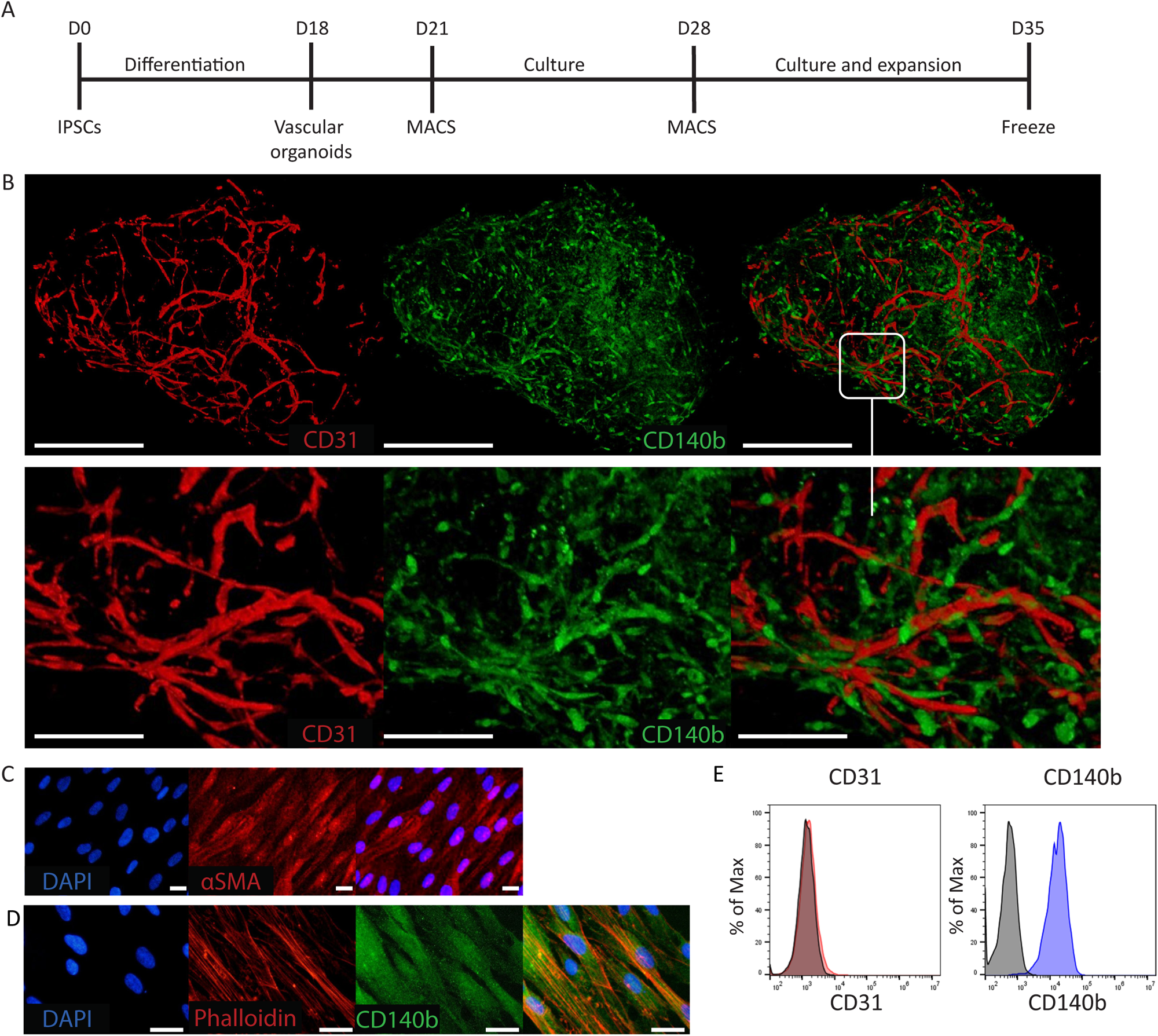

Throughout the study, human aortic VSMCs (aVSMCs) were used as a control. The phenotypic switch towards a contractile phenotype in 2D culture was induced in aVSMCs and ODMCs by a combination of low serum and TGFβ for 48 hours (T=72 h in culture, see schematics in fig. 2A). Full DMEM (with 10% serum) supplemented with both PDGFB and TGFβ was used for the synthetic phenotype, and the control groups were maintained on full DMEM (10% serum). Cell viability as measured by PrestoBlue remained unchanged in the contractile population, whereas in the control- and synthetic group this number increased significantly over time, for both aVSMCs and ODMCs (fig. 2B). In addition, gene-expression analysis of contractile markers ACTA2 and Calponin shows significant upregulation of both genes after inducing the phenotypic switch towards a contractile population, both in ODMCs and aVSMCs (fig. 2C). Immunofluorescent staining of synthetic and contractile populations (fig. 2D-E) shows similar changes in morphology and expression of Calponin protein levels in ODMCs and aVSMCs in response to phenotype induction, with for the synthetic cells showing more rhomboid shapes, and contractile cells demonstrating more cell elongation; as indicated by quantification of the aspect ratio (major axis/minor axis) (fig. 2F). Cell expression of Calponin protein was quantified by assessment of the Calponin+ area per cell (fig. 2G). The contractile populations of ODMCs and aVSMCs show comparable higher levels of Calponin+ cells, than in their respective synthetic populations. Together these data indicate that ODMCs are capable of adapting a synthetic or contractile phenotype that is induced by the different growth factors regimes, similar to primary VSMCs.

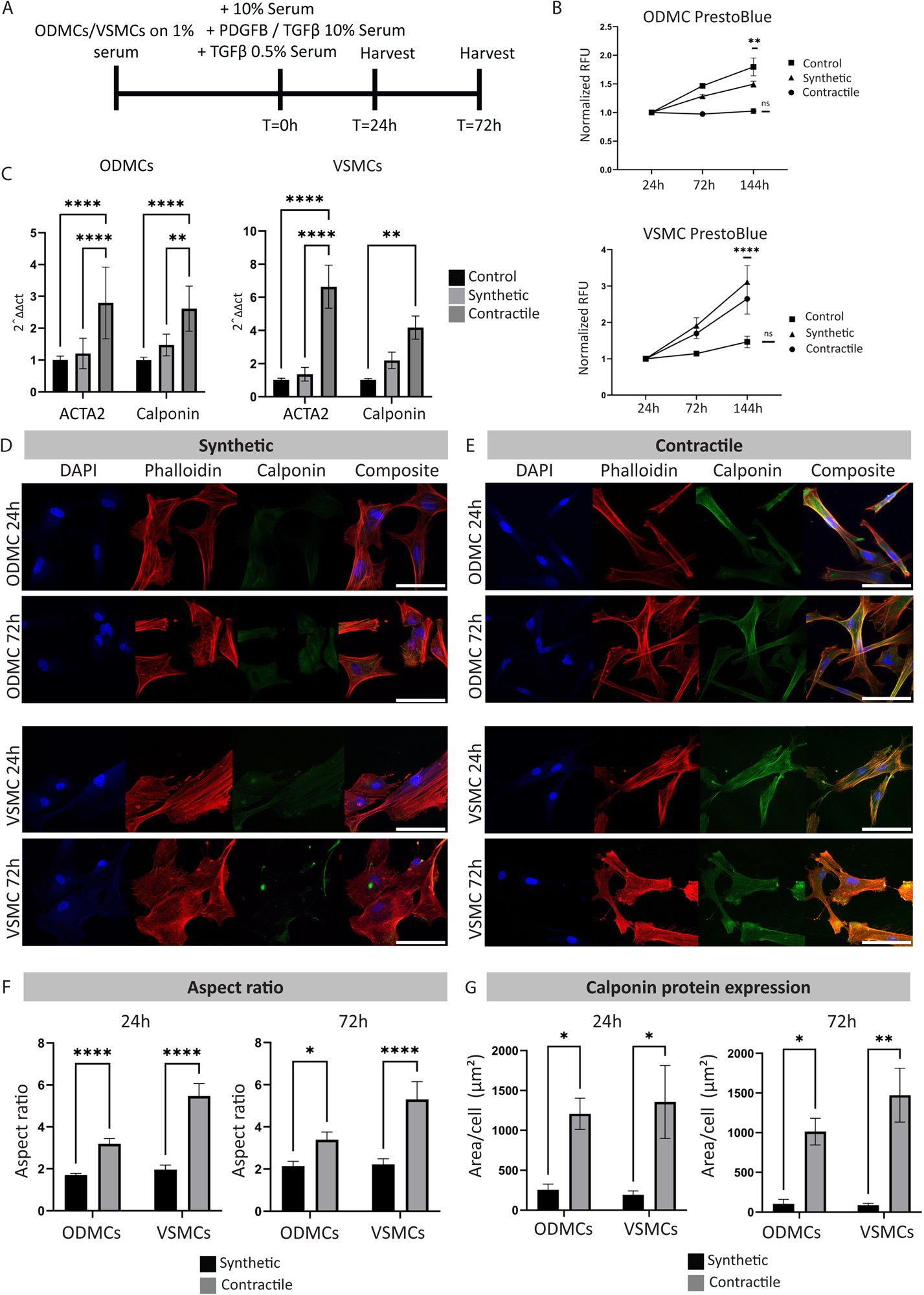

### ODMCs and VSMCs static culture in 3D in different GelMa hydrogel conditions with specific intrinsic matrix characteristics has limited impact on cell survival

Cells were seeded and statically cultured in 3D in GelMa hydrogels for a total of 72 h before harvesting (fig. 3A). Bare GelMa hydrogel characteristics were first analyzed, starting with the water absorption capacity using a swelling assay (fig. 3B). This assay tests the DOF, with higher DOF hydrogels accommodating higher degree of cross-linking, resulting in lower mass/swelling ratio (q). The q was higher for the 5% compared to the 10% gels for both DOFs (*p*<0.001 for 50 DOF, *p*<0.0001 for 80 DOF) All 80 DOF hydrogels show significant higher ratios versus 50 DOF compared to their respective percentage counterparts.

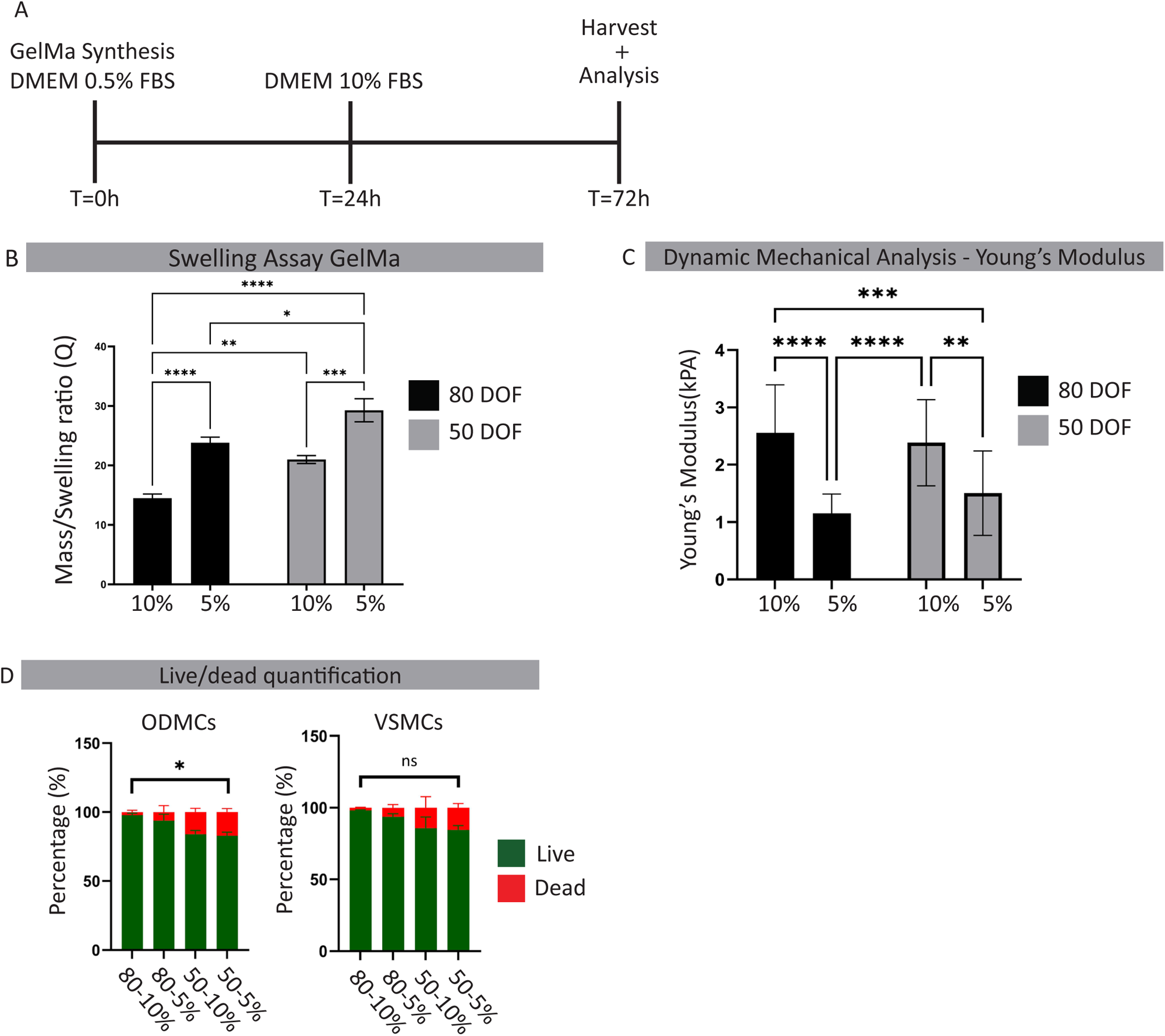

Gel mechanical characteristics were assessed by dynamic mechanical analysis (DMA) and Young’s modulus calculations are displayed as kPA (fig. 3C). The Young’s modulus reflects the material’s viscoelastic response, with higher values represent higher material resistance to deformation when subjected to mechanical forces. 10% gels demonstrate a significant higher Young’s modulus compared to the 5% gels for all DOFs (*p*<0.0001 for 80 DOF, *p*<0.01 for 50 DOF).

Calculation of the Live/dead cell ratio of ODMCs and aVSMCs in the different hydrogels after maximal 144 h of static culture show no effect of hydrogel conditions in aVSMCs, and only a significant decline in the 5% 50 DOF versus the 10% 80 DOF condition in ODMCs (Fig. 3D). Combined, these data validate the differences in intrinsic matrix properties between hydrogel conditions, with higher water absorption capacity in 50 DOF versus 80 DOF indicative of a higher crosslinking density in 80 DOF, and a higher Young’s modulus (thus higher stiffness and lower elasticity) in 10% versus 5% compositions. These conditions have limited impact on cell survival of the tested cells in static culture.

### High crosslink density and high stiffness in a static 3D GelMa environment promotes a contractile phenotype in ODMCs and aVSMCs

After 72 h of static 3D culture in the GelMa hydrogels, cells were harvested for analysis. Expression of contractile markers ACTA2, Calponin and Collagen I were assessed using qPCR and relative mRNA expression and are shown in figure 4A to evaluate the impact on cell phenotype. In ODMCs, in 5% hydrogels, 80 DOF significantly increased expression of contractile markers compared to 50 DOF (*p*<0.05 for ACTA, *p*<0.0001 for Calponin and *p*<0.01 for Collagen I). In 10% hydrogels, a similar trend was observed, but only for ACTA2, expression was significantly increased in 80 DOF versus 50 DOF (*p*<0.05) (fig. 4A). This indicates that GelMa hydrogels with higher crosslink densities (DOF) promoted adaptation to a contractile phenotype under static culture in ODMCs. For aVSMCs, a similar trend was observed with increased expression of Calponin, but also a decrease in Collagen I, in the 80 DOF-5% gels compared to 50 DOF-5% gels (*p*<0.05 for Calponin, *p*<0.01 for Collagen I). In the 10% gels, both ACTA2 and Calponin were upregulated in the 80 DOF versus 50 DOF (*p*<0.05 for ACTA, *p*<0.01 for Calponin).

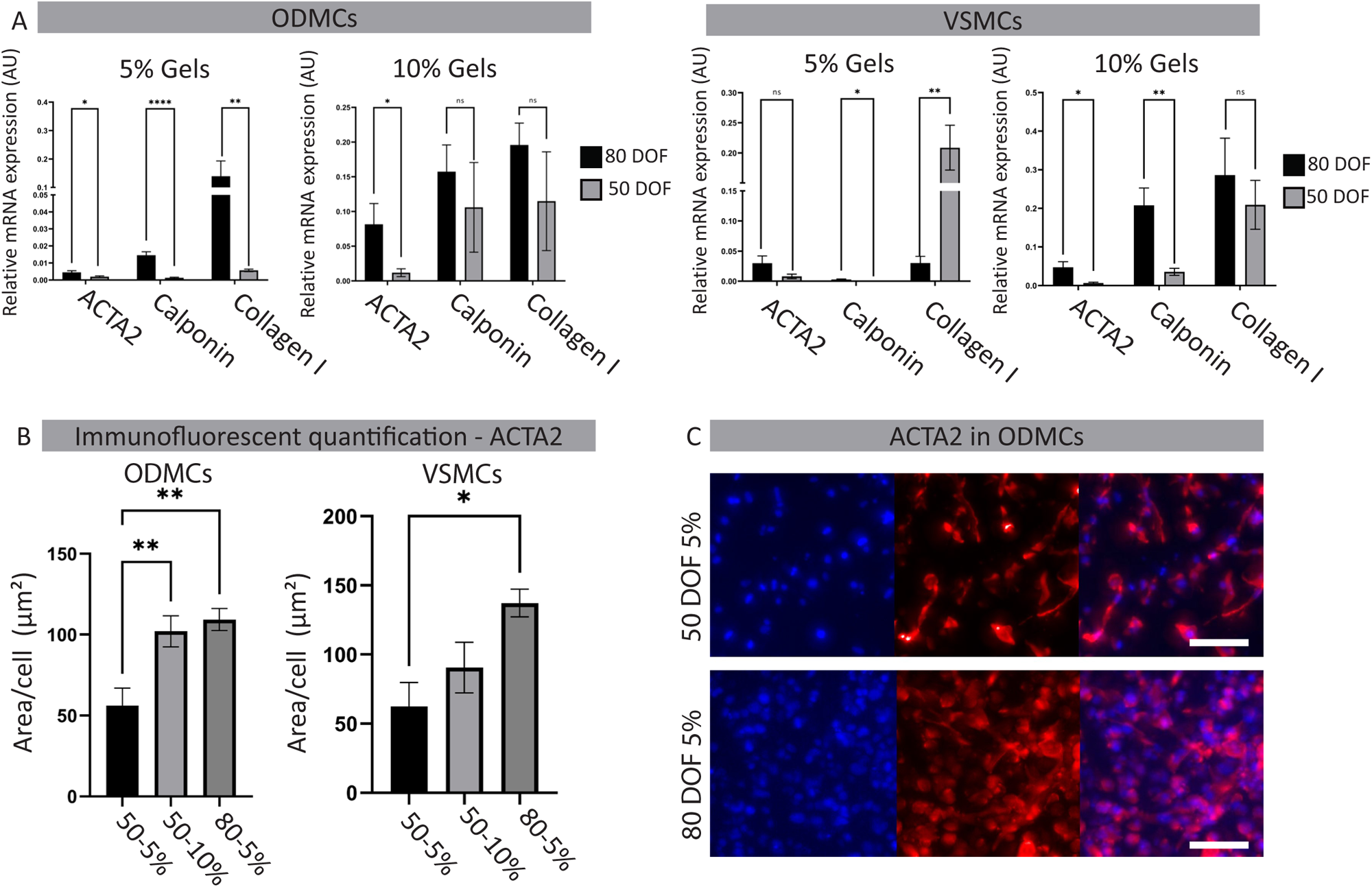

When the weight percentages of the same DOFs were compared, a significant (*p*<0.05) increased expression was observed in the ODMCs of both Calponin and Collagen I in the 10% gels compared to the 5% gels of the same DOFs (supplementary Fig 1A). For aVSMCs, there were no significant differences in expression of contractile genes in the 50 DOF conditions. In the 80 DOF, the 10% gels caused significant (*p*<0.05) upregulation of Calponin and Collagen I. These findings indicate that 3D culture in GelMa hydrogels with high young’s modulus (high stiffness, low elasticity) promotes a contractile in phenotype in ODMCs and aVSMCs.

The addition of growth factors in the different static 3D GelMa conditions did not significantly alter gene expression of contractile markers of ODMCs or aVSMCs (supplementary figure 1B). Protein levels of ACTA2 quantified by assessment of the ACTA2+ area per cell (fig. 4B) showed significant increase in 80 DOF-5% versus 50 DOF-5% gels for ODMCs and increase in 80 DOF-5% versus 50 DOF-5% in aVSMCs. Examples of the ACTA2 staining of the 50 DOF-5% and 80 DOF-5% for ODMCs are shown in figure 4C. Examples of ACTA2 stainings for all conditions for both ODMCs and aVSMCs are displayed in supplementary figure 1C. For 80 DOF-10%, both ODMCs and aVSMCs exhibited a rounded cell morphology with limited elongation, suggesting minimal interaction with the hydrogel.

### Uniaxial strain induces switch towards a contractile phenotype in 3D GelMa cultured ODMCs and aVSMCs

Using the Flexcell© Tissue train system, 10% Uniaxial strain was applied for 48 h on (72 h old) seeded gels, as displayed in the timeline in figure 5A. The effect of gel characteristics on strain patterns was assessed by comparing strain levels between the strongest (80 DOF-10% gels) and weakest (50 DOF-5%) hydrogel gels. No significant differences in strain patterns were detected (Supplementary fig. 2A). Strain analysis also reveals no significant differences in strain levels between day 0 and day 2 timepoints or between the different experiments (Supplementary fig. 2B). Gene-expression levels of contractile markers in dynamic conditions were compared to the static controls, displayed in figure 5B. 48 h strain increases the expression of contractile markers in GelMa hydrogels for multiple conditions: In ODMCs, 80 DOF-5%, 80 DOF-10%, and 50 DOF-10% GelMa hydrogels show significant increased expression of ACTA2 (*p*<0.01 for 80-10%, *p*<0.05 for 80-5% and *p*<0.01 for 50-10%) and Calponin (*p*<0.01 for 80-10%, *p*<0.05 for 80-5% and *p*<0.05 for 50-10%) after exposure to strain. For VSMCs, all conditions except the 80-5% gel showed significant increased expression of ACTA2 (*p*<0.001 for 80-10%, *p*<0.05 for 50-10% and *p*<0.001 for 50-5%). For Calponin only the cells in the 50-5% (<u>p</u><0.05) and 80-10% (*p*<0.01) gels showed significant upregulation. For collagen I, the exposure to strain did not significantly affect gene expression levels in both cell types. ACTA2 protein levels were assessed by quantification of the ACTA2+ area per cell (Fig. 5C). Strain significantly increased ACTA2+ levels in the 50 DOF gels for both cell types (*p*<0.05 for 5% gels in ODMCs, *p*<0.001 for 5% gels in VSMCs and both 10% gels).

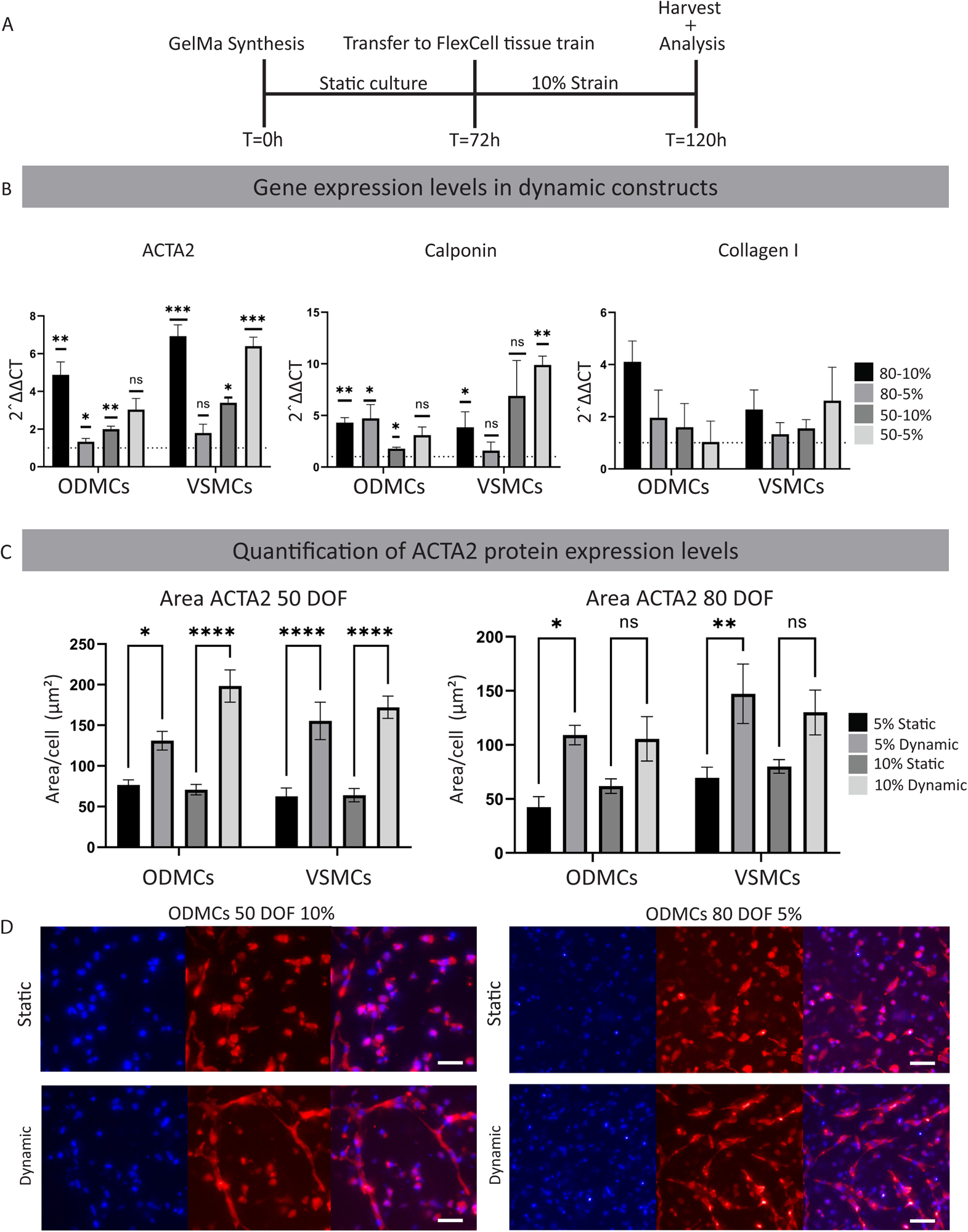

In the 80 DOF gels, there was only a significant increase in the 5% gels in both cell types (*p*<0.05 for ODMCs, *p*<0.01 for VSMCs). Examples of the ACTA2 staining of both 50-10% and 80-5% gels are shown in figure 5D. Increased elongation of both ODMCs and aVSMCs after exposure to strain is clearly visible, indicating morphological adaptation to strain. ACTA2 stainings for all conditions for both cell types are displayed in supplementary figure 3A-B.

## Discussion

Within the vascular research field, there is a notable gap in comparative studies regarding the impact of intrinsic matrix substrate properties, such as hydrogel stiffness, elasticity, and degree of crosslinking, on the cell behaviour of (h)iPSC-derived vSMCs, particularly when cultured in three-dimensional (3D) structures. Additionally, the phenotypic adaptation of (h)iPSC-derived vSMCs in response to these factors in 3D environments under cyclic strain is largely unexplored. Here, we demonstrated the ability of hiPSC ODMCs to undergo phenotype switching under different culture and 3D GelMa hydrogel conditions, similar to primary aVSMCs, illustrating their suitability of use in modelling of complex disease and potential for therapeutic interventions. The main findings are: (1) ODMCs derived from hiPSCs exhibited a VSMC phenotype, expressing key mural markers such as α-smooth muscle actin (αSMA) and CD140b. (2) ODMCs demonstrated phenotypic plasticity in response to specific culture conditions and can adopt a contractile phenotype similar to primary human VSMCs. (3) The mechanical properties in a 3D hydrogel substrate, including stiffness, elasticity, and degree of crosslinking had profound impact on both ODMCs and aVSMCs under static culture, with higher material stiffness/lower elasticity and higher crosslink density inducing a contractile phenotype. (4) Dynamic stimulation in 3D substrate using uniaxial strain further promotes a switch towards a contractile phenotype in both ODMCs and aVSMCs. Our research enhances knowledge of human iPSC-derived VSMCs, particularly ODMCs, by elucidating the influence of culture medium composition, intrinsic matrix properties, and dynamic stimuli on phenotypic changes. These findings have practical implications for tissue engineering and regenerative medicine, especially in vascular disease treatment and modeling.

### ODMCs are capable of growth-factor induced phenotype switching similar to primary human aorta derived VSMCs

ODMCs derived from vascular organoids ^30^ typically exhibit pericyte-like coverage of microcapillaries. However, when isolated and purified through CD140b sorting and cultured in VSMCs medium, they undergo a morphological transition towards VSMC-like cells. When seeded on solution electrospun vascular scaffolds, these ODMCs can form tissue structures resembling the tunica media ^29^. Notably, they form a distinct multicellular layer separate from the endothelium and contribute to the stability of these vascular grafts under flow conditions. The phenotypic characteristics of these ODMCs have not been comprehensively evaluated. Here, we demonstrated, for the first time, the phenotypic plasticity of ODMCs derived from vascular organoids.

In healthy adult vasculature, VSMCs exhibit a contractile phenotype with limited proliferation and low synthetic activity. Following vascular injury, VSMCs undergo phenotypic changes, increasing migratory, synthetic, and proliferative capacities, contributing to vascular repair but also to diseases such as atherosclerosis, cancer, and hypertension [9, 32, 33]. Notably, PDGFB induces the synthetic VSMC phenotype by downregulating contractile gene expression and promoting proliferation and migration [41, 42]. In contrast, TGF-β and Bone Morphogenetic Protein 4 (BMP4) inhibit VSMC proliferation and migration while inducing contractile gene expression [19, 45-47]. Serum deprivation enhances contractile gene expression, which can be reversed upon restoring a serum-rich medium, resulting in decreased contractile gene expression and a morphological transition of VSMCs. Notably, serum and PDGFB deprivation of human pluripotent stem cell (hPSC)-derived VSMCs have been observed to induce maturation towards a contractile phenotype [49], whereas the use of a high-serum medium in conjunction with PDGFB treatment has been found to induce the synthetic phenotype in these hPSCs.

Based on these previous reports, we used serum and PDGFB starvation, along with TGF-β treatment in our experiments to assess the capacity of ODMCs to acquire a contractile phenotype. Our results demonstrated that ODMCs, like aVSMCs, successfully acquired a contractile phenotype when exposed to the “contractile” culture conditions. This was evidenced by upregulation of contractile markers, reduction in cell proliferation rate, and elongation of cells, as compared to the control condition (ODMCs and aVSMCs in 10% serum) and the synthetic condition (ODMCs and aVSMCs in 10% serum, with PDGFB and TGF-β). Stimulation with the synthetic culture medium did not elicit any differences in marker expression, cell proliferation rate, or morphology in ODMCs or aVSMCs, compared to the control conditions. It has been observed that (prolonged) *in vitro* expansion of VSMCs can lead to a gradual loss of the contractile phenotype ^31^. The lack of response to the “synthetic” culture medium could indicate that the control conditions utilized in our experiments already maintained a more synthetic population of ODMCs and aVSMCs. Nevertheless, our findings demonstrate that ODMCs exhibit a level of phenotype plasticity that is comparable to that of aVSMCs.

### Higher crosslink density and higher stiffness in a static 3D GelMa environment promotes a contractile phenotype in ODMCs and aVSMCs

Based on gelatin modified with methacryloyl groups, GelMa is biocompatible and biodegradable and is widely used in various tissue engineering strategies, including in 3D cell printing to recapitulate blood vessels or vascularized tissues ^32^. The mechanical properties of GelMa are tunable by altering by its crosslinking conditions, including polymer concentration, degree of methacrylation, light wavelength and intensity and light exposure time ^33, 34^. The viability, function and survival of GelMa loaded cells is highly dependent on the resulting crosslink density. Here, we used two different degrees of metacrylation (or degree of functionalization, DOF), 50% and 80%. Of these two DOFs, we used two different weight percentages; 5% and 10%, to create four different hydrogels with each different intrinsic matrix properties: (1) High crosslinking, high stiffness (80 DOF 10%), (2) high crosslinking, low stiffness (80 DOF 5%), (3) low crosslinking, high stiffness (50 DOF 10%), (4) low crosslinking, low stiffness (50 DOF 5%). For vascular cells, high crosslink density in GelMa was previously shown to be detrimental to vascular network formation *in vitro* and *in vivo*, resulting in less and shorter neovessels with fewer branchpoints ^35, 36^. Although the impact of a high degree of GelMa crosslinking on VSMCs was not investigated, the mesenchymal stem cells that were used for vascular support in these studies showed significant reduction in perivascular recruitment by neovessels and *in situ* impairment of differentiation into mural cells. A higher degree of crosslinking has also been associated with reduced cell spreading capacity by increasing the physical matrix barrier and reduction in pore size ^37^. In line with these observations, ODMCs and aVSMCs in 80 DOF 10% hydrogels showed limited elongation in cell morphology compared to cells in 50 DOF (10% and 5%) or 80 DOF 5% under static conditions, indicative of impairment in cell spreading. Dynamic stimulation of the 80 DOF 10% condition only induced limited morphological adaptation in aVSMCs but not ODMCs, and strained aVSMCs displayed non-typical cell thinning or enlargement instead of elongation (Supplementary figure 2). A higher degree crosslinking of GelMa was previously reported to reduce expression of mural cell markers of mesenchymal stem cells ^35^, but the impact on VSMC phenotype switching in hiPSC-derived mural cells remained to be invested. Here we observed under static conditions a significant higher expression of contractile markers in 80 DOF versus 50 DOF, in 5% and to a lesser extend in 10% hydrogels, with ODMCs performing better than aVSMCs (Fig 4A, B), demonstrating that higher crosslinking of 3D hydrogels promotes a contractile phenotype.

Phenotype determination in primary vSMCs may also be controlled by hydrogel stiffness. The limited available data on the effect of substrate stiffness on VSMC behavior is derived from 2D experiments. Higher substrate stiffness in these 2D studies have highlighted the impact of different ECM component with e.g. collagen type I coating on increasingly stiffer substrates reducing (synthetic phenotype associated) VSMCs migration, whereas higher substrate stiffness with fibronectin coating promoted migratory behaviour ^24^. Notably, it has been indicated that the migratory response substrate stiffness within a range of 1.0 to 308 kPa (Young’s modulus) is biphasic, implying that there is an optimal substrate stiffness for maximal migration ^25^. Another interesting observation is that substrates with higher stiffness require lower density of ECM (fibronectin) coating to achieve a similar migratory response in VSMCs than substrates with lower stiffness ^25^. These findings indicate that the response of VSMCs to substrate stiffness in 2D is highly dependent on the assessed stiffness range and ECM component type and density. How these findings will translate in a more physiologically relevant 3D environment, in particular for hiPSC-derived VSMCs, remains largely underexplored. A recent comparative investigation focusing on the cyclic stretching stimulation of human VSMCs showed contrasting outcomes between 2D and 3D models, with contractile protein expression remaining unaltered under 2D stretching conditions, and exhibiting a notable increase within 3D collagen matrix conditions ^28^. These disparities underscore the possible critical influence of extracellular dimensionality (2D or 3D) on cellular responses to mechanical stimulation, emphasizing the urgent need to broaden the scope of current investigations in this domain. By comparing 5% with 10% hydrogels, our data showed that higher matrix stiffness in 3D GelMa hydrogel under static conditions increased expression of contractile markers in ODMCs in both 80 and 50 DOF conditions. The same effect was observed to a lesser extend for aVSMCs for 80 DOF gels (Supplementary figure 1C, figure 4B). These results are partially in line with previous findings: Peyton et al. showed that adjusting the modulus within the range of 0.45 to 5.8 kPa resulted in modulation of cytoskeletal assembly in in human primary VSCMs in a 3D PEG-fibrinogen based static hydrogel, with stiff matrices exhibited a slightly elevated level of F-actin bundling ^38^. However, the expression of contractile markers in Peyton’s study was only increased in response to higher matrix stiffness after constitutive RhoA activation, which may be attributed to the use of different hydrogels (GelMa versus PEG-fibrinogen). Similar to Peyton’s findings, static 3D culture with different matrix stiffness (5% versus 10% gels) had no effect on cell survival of ODMCs and aVSMCs. Crosslinking density had also no impact on the survival of ODMCs (Figure 3D). Combined, these findings demonstrate, for the first time, that increased stiffness in GelMa based hydrogels promote a viable contractile phenotype in hiPSC-derived VSMCs.

### Uniaxial strain induces phenotypic switch towards a contractile population of smooth muscle cells in ODMCs and aVSMCs

The effect of cyclic strain on the morphology and function of VSMCs has been described predominantly in 2D setups. The use of cyclic strain in 2D within the pathologically relevant range (>15%) has been shown to induce DNA synthesis through increased reactive oxygen species (ROS) production and NF-kB pathway activation ^39^. Conversely, physiological strain levels (10%) inhibit VSMC proliferation by upregulating p21 expression and promoting apoptosis ^40, 41^. For human iPSC-derived VSMCs, data on cyclic strain is very limited, with findings that indicate an ability to cytoskeletal remodeling to 2D strain similar to primary VSMCs in a progeria-on-a-chip model ^42^. In relation to expression of phenotypic markers, up and down regulation of contractile markers have been reported following VSMC exposure to cyclic strain ^43–47^. Most notably, Bono et al. (2016) compared the impact of cyclic strain on VSMCs cultured in type I collagen substrate in both 2D and 3D environments ^28^. In the 2D model, they reported a downregulation of contractile proteins (αSMA and Calponin) in the strained versus static samples. In contrast, a twofold increase in αSMA and 14-fold increase in Calponin expression was observed in the 3D conditions when exposed to cyclic stain. This coincided with a difference in morphological adaptation with a perpendicular (80-90 degrees) alignment in the 2D cultured versus parallel (0-10 degrees) alignment in 3D cultured VSMCs, in relation to the strain direction. These findings imply that VSMCs adaptation to cyclic strain is profoundly different in 2D versus 3D conditions. In line with these 3D findings from Bono’s study, we observed that cyclic strain significantly increased the expression of contractile markers and caused elongation of ODMCs and VSMCs in direction of the applied strain in all our four 3D hydrogel conditions. Crosslink density (80 DOF and 50 DOF) and matrix stiffness/elasticity (10% and 5%) did not affect this contractile switch in both cell types. Strain pattern comparison of the 80 DOF 10% versus the 50 DOF 5% gel did not show any significant differences, indicating variations within this range may have limited impact on the local strain levels of what the cells experience on an individual level. Future research should investigate hydrogels with a higher range in crosslinking and stiffness to assess the impact of hydrogel properties on the conveyance of strain from the tissue to the cellular level.

In this study, we demonstrate the phenotypic plasticity of hiPSC-derived ODMCs, which have the capacity to adopt a contractile phenotype in response to growth factor stimulation in 2D. In addition, 3D culture in GelMa hydrogels under static condition showed that higher material stiffness, lower elasticity, and higher crosslink density induced a contractile phenotype in these cells, similar to aVSMCs. Dynamic stimulation in the 3D substrate using uniaxial strain further promoted a switch towards a contractile phenotype in both ODMCs and primary VSMCs.

These findings underscore the significance of optimizing matrix properties within a (dynamic) 3D environment, and contribute to the advancement of sophisticated human disease models and vascular tissue engineering strategies.

## Acknowledgements

This work was funded by the REGMEDXB cardiovascular moonshot consortium and the NWO Vidi grant (no. 91714302 to CC). The authors gratefully acknowledge the Gravitation Program “Materials Driven Regeneration”, funded by the Netherlands Organization for Scientific Research (024.003.013).

## Author Contributions

**EMM** wrote the manuscript together with **RG** and **CC.** Experiments were executed by **EMM, RG, CVD, RM, IC, TBW** and **YA. HC** performed strain analysis**. AIPMS, MCV** and **CC** supervised the project. All authors read, revised and accepted the manuscript.

